# NanoString Technologies Neuropathology Panel Produces Unreliable Measurements of Mouse Hippocampal Gene Expression

**DOI:** 10.1101/2025.07.24.666661

**Authors:** Justin S. Rhodes, Meghan G. Connolly

**Affiliations:** Department of Psychology, University of Illinois, Urbana-Champaign, IL, USA; Neuroscience Program, University of Illinois, Urbana-Champaign, IL, USA; Department of Chemistry, University of Alberta, Edmonton, AB, Canada

**Author notes:** **Corresponding author:** Justin S. Rhodes, Department of Psychology, 603 East Daniel St., Champaign, IL 61820.

**Keywords:** Nanostring, nCounter, gene expression, RNA, multiplex

## Abstract

Technologies for measuring gene expression (i.e., the number of RNA transcripts) of large numbers of genes simultaneously in specific tissues have exploded in recent years. Current methods include high-throughput RNA sequencing (RNA-seq), transcript counting platforms like NanoString’s nCounter®, and spatially resolved techniques based on fluorescent in situ hybridization (FISH). Several studies have evaluated the reliability of these different methods and performance in comparison to one another. Typically, technical reliability, as measured by Pearson’s correlation of two measurements of the same sample, is usually well above 90%, and is statistically significant even for small sample sizes (e.g., 8 samples measured twice). We performed an experiment where we aimed to compare hippocampal gene expression between 3 groups (n=5 per group) of young adult male C57BL/6J mice. Before sampling, the groups were treated with either repeated injections of PBS (vehicle), extracellular vesicles taken from the blood plasma of sedentary mice (SedVs) or exercising mice (ExerVs). The hippocampus was dissected, and RNA purified using standard methods. The samples were analyzed using the NanoString Neuropathology panel, that measures 770 genes simultaneously. To estimate reliability, we measured 8 of the samples twice in two separate assays. Surprisingly, only 85 genes showed a significant Pearson’s correlation (p<0.05), and none of these met false discovery significance (all q<0.05). To confirm that no errors were made transferring labels, the individual samples were permuted to see whether a different assignment could recover a greater number of positive correlations. Results showed that the original assignment was best suggesting no errors in sample assignments were made. We conclude that the Nanostring neuropathology panel produces unreliable data for mouse hippocampal gene expression.

## Introduction

High throughput technologies for measuring gene expression (i.e., the amount of mRNA transcripts) in biological tissues have rapidly advanced in recent years. Examples include single cell or single nuclei RNA sequencing, bulk RNA sequencing, spatial transcriptomics (e.g., Visium HD, Xenium), MERFISH, among many others. All these methods are capable of measuring hundreds to thousands of genes at a time in isolated tissues with high accuracy, precision, and reliability (Stark, Grzelak, & Hadfield, 2019; Stuart & Satija, 2019).

Recently a new method has emerged, the NanoString’s nCounter. In brief, it uses two sequence-specific probes—a fluorescently labeled reporter probe and a biotin-labeled capture probe—that hybridize to each target RNA. After hybridization and purification, the system digitally counts each unique fluorescent barcode, providing a direct count of RNA molecules. The approach is supposed to provide high sensitivity, precision, and reproducibility. It is also sometimes preferred since it does not measure the entire transcriptome, but rather is a more targeted approach focused on a large number of genes that have already been annotated and are relevant for the biological question. This was supposed to be the case for the NanoString Technologies Neuropathology Panel. On the website, it says the genes were chosen because they are, “…involved in six fundamental themes of neurodegeneration: neurotransmission, neuron-glia interaction, neuroplasticity, cell structure integrity, neuroinflammation, and metabolism.”

When measuring gene expression using any of these methods, there are always at least two sources of variation. One source is individual variation, which comes from possible genetic and/or environmental differences (e.g., treatment groups, genotypes, etc.) between individual samples. The other source is technical variation, which is determined by the precision of the measuring device. Technical variation is commonly estimated by measuring the same sample twice and calculating the Pearson’s correlation between the two measurements. Typically, for gene expression, the correlation is above 90%. It is obviously important to keep technical variation low, because the more measurement error, the more difficult it will be to detect the biological effects of interest (e.g., the genetic or treatment effects) (Schurch et al., 2016).

We used NanoString Technologies Neuropathology Panel to measure gene expression in the hippocampus of 15 mice. In 8 of these mice, we measured gene expression twice in two separate NanoString Technologies Neuropathology Panels to measure technical reliability. The purpose of this paper is to report on the technical reliability of the NanoString Technologies Neuropathology Panel. Thus, the details of the experiment’s aim and design are not pertinent. Nevertheless, for the sake of context, our aim was to see how extracellular vesicles (EVs) from exercising mice (ExerVs) would affect hippocampal gene expression in sedentary mice. In the same mice that were measured for gene expression reported herein, half the hippocampus was processed to measure adult hippocampal neurogenesis using the BrdU method and we observed ExerVs increased neurogenesis relative to controls (paper under review). The effect was replicated in a second cohort. Thus, we wanted to see how the ExerVs may have altered gene expression in the hippocampus in parallel with the increased neurogenesis to better understand how the ExerVs may have contributed to increased neurogenesis.

## Materials and Methods

### Experimental subjects

A total of 15 adult male C57BL/6J mice (4-month-old) were group-housed and maintained in ventilated cages with ad libitum access to food and water throughout the experiments. The colony room was maintained on a 12-hour light/dark cycle. The experiments were approved by the University of Illinois Urbana-Champaign Institutional Animal Care and Use Committee (IACUC) (#22062) and followed the Guide for the Care and Use of Laboratory Animals. For details on the experimental treatments, please see (Connolly et al., 2025). The 15 mice were divided into three groups. One was treated with PBS injections, one with extracellular vesicles (EVs) from exercising mice (ExerVs), and one with EVs from sedentary mice (SedVs) to serve as another control in addition to PBS.

### Sample collection

Mice were anesthetized with an ip injection of Fatal Plus (sodium pentobarbital) solution, then euthanized by transcardial perfusion of ice cold 0.9% saline. The brains were immediately removed and divided in half. One half was used for BrdU immunohistochemistry to measure neurogenesis (results are not reported here). The hippocampus was rapidly dissected from the remaining hemisphere and snap-frozen for RNA extraction and gene expression analysis.

### RNA extraction

RNA from the hippocampus was extracted using the Qiagen RNeasy Lipid Tissue Mini Kit (Qiagen, Cat# 74804) by following the manufacturer directions. Briefly, tissue was homogenized using a motorized pestle in 0.6 ml of the Qiazol Lysis Reagent. The homogenate was then incubated for five minutes and then 0.2 ml of chloroform was added to promote phase separation of RNA. This mixture was then centrifuged and the top layer formed during centrifugation was collected. 70% ethanol was added to the collected layer to precipitate the RNA. The mixture was then moved to a spin column tube and centrifuged allowing the liquid to pass through tube and the RNA to bind to the clear silica-based membrane contained in the cartridge tube. Any impurities in the RNA sample were removed through multiple wash steps. The purified, bound RNA was then eluted in RNase-free water. Immediately following this process, the samples were assessed for purity (260/280 ratio) and concentration using a Take3 plate read on a Gen5 Epoch spectrophotometer (BioTek Instruments, Highlands Park, VT). The purified RNA was stored at −80°C until use for the NanoString Assay.

### NanoString assay

Total RNA was isolated from 15 samples and submitted to the Tumor Engineering and Phenotyping (TEP) facility at the University of Illinois Urbana-Champaign. RNA integrity was confirmed by gel electrophoresis, and samples were assayed on the nCounter Neuropathology Panel (NanoString Technologies, Seattle, WA) according to the manufacturer’s protocol. Of the 15 samples, 8 were run in technical replicate to assess reproducibility across runs. Raw NanoString data were imported into nSolver□4.0 software and processed using the recommended workflow, including system- and imaging-QC flag inspection, background subtraction using negative controls, and normalization using positive spike-in controls and geometric-mean-adjusted housekeeping genes.

### Statistical analysis

All statistics were performed in R (version 4.4). Pearson’s correlations were calculated to assess the consistency between the two repeated measurements within the eight samples, for each of the 770 measured genes. To verify that no errors in sample assignments were made, the individual samples were permuted to see whether a different assignment could recover a greater number of positive correlations. For each permuted assignment, the number of significant positive correlations detected across all 770 genes was recorded. This number represents the number of genes that could be reliably measured. The number of reliably measured genes for the permuted assignments was then plotted in a histogram to compare to the number for the actual assignment. See Supplementary file 1 for the R code to implement the analysis.

## Results

### RNA quality

RNA purity was high across all samples, with 260/280 ratios ranging from 2.129 to 2.169 (mean ± SD = 2.147 ± 0.010). RNA concentrations ranged from 238.156 to 700.254 ng/µL.

### Reliability

Raw output from the NanoString Technologies Neuropathology Panel is available as a supplementary file for users to explore (Supplementary box folder: https://uofi.box.com/s/d257iaq35qukjhr71k1lgxycowxbs5ho). Surprisingly, only 85 genes showed a significant Pearson’s correlation (p<0.05), and none of these met false discovery significance (all q>0.05) (see Supplementary Table 1). The permutation analysis showed that the original assignment had the highest number of reliably measured genes thus suggesting no errors in sample assignments were made (Figure 1).

**Table 1.** RNA purity and concentration for all samples submitted for NanoString analysis. Each sample’s treatment group, 260/280 absorbance ratio (as a measure of RNA purity), and RNA concentration (ng/µL) are shown. RNA integrity was confirmed prior to analysis.

| Sample | Treatment | 260/280 | ng/μL conc. |
| --- | --- | --- | --- |
| 1 | PBS | 2.137 | 238.156 |
| 2 | PBS | 2.169 | 523.198 |
| 3 | PBS | 2.137 | 314.826 |
| 4 | PBS | 2.163 | 700.254 |
| 5 | PBS | 2.153 | 564.807 |
| 6 | SedV | 2.137 | 249.335 |
| 7 | SedV | 2.156 | 412.248 |
| 8 | SedV | 2.156 | 314.224 |
| 9 | SedV | 2.13 | 339.589 |
| 10 | SedV | 2.157 | 422.073 |
| 11 | ExerV | 2.15 | 477.251 |
| 12 | ExerV | 2.129 | 341.312 |
| 13 | ExerV | 2.141 | 374.91 |
| 14 | ExerV | 2.152 | 462.752 |
| 15 | ExerV | 2.153 | 353.933 |

**Figure 1.**
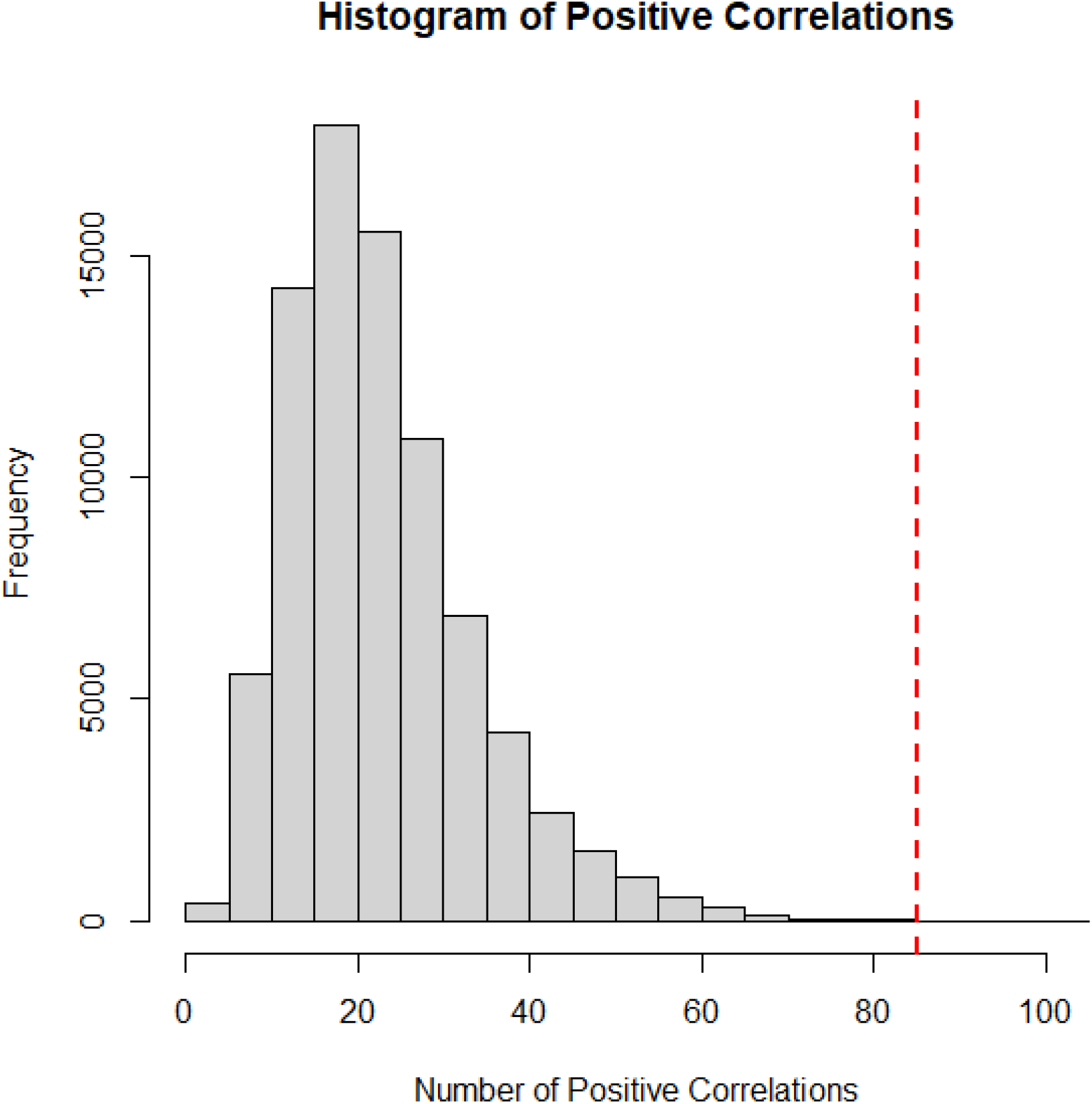
Histogram showing the results of a permutation analysis in which sample identities were randomly reassigned across iterations. For each permutation, Pearson’s correlations were computed for all 770 genes in the NanoString neuropathology panel. The histogram displays the number of significant positive correlations (p < 0.05) detected in each iteration. The red horizontal dashed line represents the number of significant correlations obtained using the original, assumed-correct sample assignments. This value lies at the extreme upper end of the distribution, providing strong evidence that the sample IDs were correctly assigned.

## Discussion

We conclude that the NanoString Neuropathology Panel produced unreliable gene expression data in mouse hippocampal tissue under the conditions tested. Despite using high-quality RNA, following the manufacturer’s protocol, and selecting a panel specifically developed for use with mouse brain tissue, only 85 of the 770 genes showed significant correlations across technical replicates, and none survived correction for multiple comparisons. Given that the Neuropathology Panel is explicitly marketed for use in mouse brain samples, these results point to a deeper issue with the technical performance of the platform itself. This degree of variability is unexpected and unacceptable for quantitative transcriptomic analysis, especially when compared to the high reproducibility typically observed in RNA-seq or even other multiplexed assays. Our findings suggest that, at least in our hands, the NanoString nCounter system does not provide sufficient technical reliability to support meaningful comparisons of hippocampal gene expression in mice. These results raise concerns about the broader utility of the Neuropathology Panel for experimental applications in neuroscience and highlight the importance of technical validation, even when using commercially available tools marketed for specific tissues and species.

